# Dead infant carrying by chimpanzee mothers in the Budongo Forest

**DOI:** 10.1101/2021.12.22.473786

**Authors:** Adrian Soldati, Pawel Fedurek, Catherine Crockford, Sam Adue, John Walter Akankwasa, Caroline Asiimwe, Jackson Asua, Gideon Atayo, Boscou Chandia, Elodie Freymann, Caroline Fryns, Geresomu Muhumuza, Derry Taylor, Klaus Zuberbühler, Catherine Hobaiter

## Abstract

It has been suggested that non-human primates can respond to deceased conspecifics in ways that suggest they experience psychological states not unlike humans, some of which could indicate they exhibit a notion of death. Here, we report long-term demographic data from two East African chimpanzee groups. During a combined 40-year observation period we recorded 191 births of which 68 died in infancy, mostly within the first year. We documented the post-mortem behaviour of the mothers and describe nine occasions where Budongo chimpanzee mothers carried infants for 1-3 days after their death, usually until the body started to decompose. We also observed three additional cases of extended carrying lasting for more than two weeks, one of which was followed by the unusual extended carrying of an object and another which lasted three months. In each case, the corpses mummified. In addition, we report four instances of recurring dead infant carrying by mothers, three of whom carried the corpse for longer during the second instance. We discuss these observations in view of functional hypotheses of dead infant carrying in primates and the potential proximate mechanisms involved in this behaviour.

## INTRODUCTION

Primate thanatology, the study of the behaviour and underlying physiological and psychological factors associated with dead or dying individuals in non-human primates (hereafter primates), continues to raise important questions about human uniqueness (Anderson 2011, 2018; Anderson et al. 2018). Humans do not immediately abandon their dead but exhibit a plethora of post-mortem social behaviour towards them (Parkes et al. 1997). These activities can last for days or weeks but are typically terminated by the onset of physical decay, although certain cultures continue to interact with the deceased for long after (e.g., Hollan 1995). Humans experiencing the loss of socially close individuals undergo significant psychological trauma with long-term physiological effects, including symptoms of post-traumatic stress disorder, anxiety and depression (Figley et al. 1997; Parkes et al. 1997; Lannen et al. 2008). There is archaeological evidence that tending the dead evolved before modern humans (Martinón-Torres et al. 2021), with mortuary behaviour documented in *Homo sapiens neanderthalensis* (Rendu et al. 2014) and *Homo naledi* (Dirks et al. 2015), having been interpreted as an indication of some abstract notion of death and understanding of irreversible loss (Pettitt 2018). Given our biological and social similarities, other primate species – in particular other great apes – may experience similar cognitive and physiological changes. Cross-species comparisons, especially in primates, are often used to reveal past evolutionary trajectories of the hominid lineage. Primate behaviour and physiology in relation to deceased individuals provide valuable data to better understand the origins of why humans are so powerfully affected by death. More generally, primate responses to death may provide further insight into other aspects of animal cognition, such as animacy or the perception of time (Gonçalves and Carvalho 2019).

Because death is unpredictable and rarely observed in wild primates, the available datasets are usually anecdotal, and descriptions of events are often incomplete (Watson and Matsuzawa 2018; Ramsay and Teichroeb 2019). Nevertheless, an increasing number of primate groups have been habituated to human observers (Kappeler et al. 2012), which has led to more frequent reporting and more systematic efforts to extract patterns of behaviour in responses to death (Anderson 2020). These observations have led to claims that primates respond to death in ways that are similar to humans, by producing strong emotional, social, and exploratory responses (Watson and Matsuzawa 2018; Gonçalves and Carvalho 2019). Among non-human animals, the emphasis on primates may result from easier detection or the relatively large number of long-term studies. However, observations from corvid, elephant, and cetacean species suggest that attentive responses towards dead conspecifics may be widespread among long-lived highly-social species (Reggente et al. 2016; Gonçalves and Biro 2018; Bercovitch 2020).

Here, we focus on a particularly remarkable behaviour seen in many primates, dead infant carrying by mothers, which in chimpanzees typically occurs for a period of up to three days (Gonçalves and Carvalho 2019). Dead infant carrying (also referred as infant corpse carrying) is the most frequently reported thanatological behaviour and shows substantial variation in how it is expressed across primate species (Fernández-Fueyo et al. 2021; Watson and Matsuzawa 2018). In addition to chimpanzees (Matsuzawa 1997; Hosaka et al. 2000; Kooriyama 2009; Biro et al. 2010), dead infant carrying has been reported in bonobos (Fowler and Hohmann 2010; Tokuyama et al. 2017), gorillas (Warren and Williamson 2004; Masi 2020), chacma baboons (Carter et al. 2020), red colobus (Georgiev et al. 2019), geladas (Fashing et al. 2011), bonnet and lion-tailed macaques (Das et al. 2019), Japanese macaques (Sugiyama et al. 2009; Takeshita et al. 2020), and vervet monkeys (Botting and van de Waal 2020) among others (Fernández-Fueyo et al. 2021), while failed apparent attempts at carrying have been observed in ring-tailed lemurs (Nakamichi et al. 1996) and marmosets (Thompson et al. 2020). Several hypotheses have been put forward in an attempt to explain the function of and motivation for this puzzling behaviour (for a review see: Watson and Matsuzawa 2018; Gonçalves and Carvalho 2019). We summarise six of the main current hypotheses from the literature to provide a framework for evaluating the potential functions and mechanisms of new observations and discuss their implications.

A first group of hypotheses presumes that primate mothers are unable to understand the ramifications of death and their behavioural responses are side effects of evolved physiological mechanisms. The ‘unawareness hypothesis’ states that mothers are unable to discriminate between temporarily unresponsive and irreversibly deceased individuals and continue providing maternal care (e.g., grooming) and trying to elicit responsiveness (e.g., poking, smelling) in order to avoid the costs of premature abandonment (Hrdy 1999). Given that the decomposition of the corpse can be mediated by local climate, dry and particularly hot or cold conditions favouring mummification should also favour prolonged carrying of dead infants (‘climate hypothesis’: Matsuzawa 1997; Fashing et al. 2011). The ‘post-parturient condition’ hypothesis (also referred as ‘hormonal’, see Gonçalves and Carvalho 2019) proposes that the maternal physiological conditions associated with pregnancy and birth favour persistent care of the dead infant as long as the mother is lactating or until resumption of ovulation (Biro et al. 2010; Kaplan 1973). After giving birth, the endocrine system of the mother releases hormones (e.g., oxytocin) that stimulate maternal behaviours (Keverne 1988; Bercovitch 2020). Thus, when mothers lose their infants close to birth, they are expected to continue to invest in infant care, increasing the likelihood and duration of post-mortem behaviour.

A second kind of hypothesis assumes that primate mothers can have a notion of death, provided they have relevant personal experience. The ‘learning about death’ hypothesis states that chimpanzees do not intuitively understand death but can learn the notion by attending to relevant cues (Cronin et al. 2011). Here, the predication is that younger or inexperienced mothers are more likely to carry dead-infants to learn about death (Watson and Matsuzawa 2018). Specifically, experiencing multiple infant deaths (their own or others’) and interacting with more living infants and/or for longer periods of time might increase mothers’ awareness of the irreversible change of death. The ‘grief-management hypothesis’ assumes that chimpanzees possess a notion of death and suggests dead-infant carrying represents a strategy to cope with grief or the stress associated with infant loss (e.g., Takeshita et al. 2020). This hypothesis predicts that mothers able to carry their dead infants experience lower levels of ‘stress’ hormones (i.e., glucocorticoids) (Nicolson 1991).

A third group of hypotheses are agnostic about whether chimpanzees possess a notion of death but propose various adaptive mechanisms that favour post-mortem mothering behaviour. The ‘learning-to-mother hypothesis’ states that dead-infant carrying improves maternal skills (Warren and Williamson 2004), predicting that the behaviour should mainly be observed in inexperienced primiparous females. The ‘maternal-bond strength hypothesis’ predicts that mothers with older infants share a stronger bond, due to more extensive association and interactions between them, and are more likely to show extensive carrying (Watson and Matsuzawa 2018).

The existence of multiple – sometime contrasting – hypotheses is likely a reflection of both the small and highly variable data available and the diversity of potential drivers of this behaviour across different species and individuals. A recent systematic study on 18 primate species (n=48 cases) showed that duration of infant carrying is affected by the age of the mother, with older mothers carrying for longer periods (Das et al. 2019). However, some species do not fit this pattern, with previous findings by Sugiyama and colleagues (2009) detecting no effect of age in a large study on Japanese macaques. Infants that died of sickness were carried for longer than those who were stillborn or victims of infanticide, but the age of the infant did not influence duration of carrying (Das et al. 2019). A study using the largest primate database to date (n=50 species and n=409 cases) found that dead infant carrying was more likely to occur when the cause of death was non-traumatic, when mothers were younger, and that older infants were carried for shorter periods (Fernández-Fueyo et al. 2021). While dead infant carrying is a shared behaviour in primates, the frequency of occurrence, duration of carrying, and ease of observation, vary considerably between species and individuals. Nevertheless, given the observations currently available, carrying duration seems to be the longest in great apes, particularly in chimpanzees (Das et al. 2019).

Chimpanzee mothers typically carry their dead infants for a few days, though a recent analysis of the largest chimpanzee dataset (n=33 cases) did not provide clear support for any of the previous hypotheses (Lonsdorf et al. 2020). Lonsdorf and colleagues (2020) proposed that the ‘unawareness hypothesis’ is unlikely because of the presence of atypical carrying postures and sensory cues after death as compared to those displayed towards living infants. However, while a decomposing body may be perceived differently from a living one through several sensory modalities (Gonçalves and Biro 2018), chimpanzee mothers may also be responding to an inability of their infant to grasp, rather than a recognition of its status as living or dead. Here, we report observations of dead infant carrying by female chimpanzees using a 40-year dataset of two study groups of the Budongo Forest, Uganda, including three detailed observations of extended dead infant carrying by two different females.

## METHODS

### Study site and subjects

The Budongo Forest Reserve is a semi-deciduous tropical rain forest located along the Western Rift Valley in Uganda. This reserve is made of 793 km^2^ of protected forest and grassland, including 482 km^2^ of continuous forest cover (Eggeling 1947) . The reserve contains a population of approximated 600 East African chimpanzees (*Pan troglodytes schweinfurthii*). Our observations took place in two adjacent communities, Sonso and Waibira, studied regularly by researchers and followed on a daily basis by field assistants since 1990 and 2011 respectively, contributing a combined 43-years of long-term observational records (Reynolds 2005; Samuni et al. 2014).

At the time of the events, the Waibira community contained an estimated 120 individuals, 96 of which could be individually recognised. Individuals involved in the first event were Ketie (KET), a 20-year old adult primiparous female (estimated birth 1998) and her 2-year old female infant Karyo (KYO) born in December 2015. The Sonso community contained 65 named individuals, in addition to three unnamed females in the process of immigrating. The individuals involved in the event were Upesi (UP), a 21-year old parous female (estimated birth 1999) and a) in the second event, her recently born unnamed unsexed infant UP3, born mid-September 2020 and b) in the third event, her fourth born unnamed unsexed infant UP4, estimated birth 7th August 2021. Her first two infants (born in 2017 and 2018) were victims of within-community infanticide before reaching a month old (see Leroux et al. 2021 for one reported case).

We considered the scope for bias in our study subjects by using the STRANGE framework to report potential sampling biases in our study (Webster and Rutz 2020; Rutz and Webster 2021). The Sonso community are of typical size whereas the Waibira community are particularly large as compared to that of other chimpanzees (in a recent comparison of 18 groups across three subspecies: *P.t. schweinfurthii, P.t. troglodytes, P.t. verus*; communities range from 7-144 individuals with a mean 42; within these data the East African sub-species (*P.t. schweinfurthii*) range is 18-144 with a median 49; Wilson et al. 2014). Sonso have a typical female-biased sex ratio among mature individuals (M:F; 1:1.7), whereas the Waibira community have more unusual evenly-balanced sex ratio among mature individuals (M:F; 1:1.2; mean among 9 *P.t. schweinfurthii* communities 1:1.7; Wilson et al. 2014). Of relevance to sampling biases in infant mortality and opportunities to carry dead infants, the Sonso community experience high levels of infanticide (Lowe et al. 2019, 2020). Characterised by a medium altitude rainforest (∼1100 m) with significant annual rainfall (∼1500 mm per year), the area is slightly more seasonal than true rainforest with a distinct dry-season during December-March and a drier season during June-August (Newton-Fisher 1999).

Observations of wild chimpanzees in dense secondary forest are often challenging and individuals in highly fission-fusion East African chimpanzee communities may not be observed for days or weeks (Badihi et al. Preprint). In addition, we often choose not to follow, or only to follow at an extended distance, individuals who are experiencing high-stress events, such as maternal loss. We considered a mother to be carrying a dead infant if they were seen to be doing so on the day following their infant’s death. In doing so, we may underestimate occurrences of shorter carries (i.e., under a day), or the total duration of carries. Similarly, as we rarely observed the moment that the mother stopped carrying the dead infant, our estimation of duration may be particularly conservative. However, given the challenges in discriminating the precise moment of infant death from for example unconsciousness, we felt that a conservative approach was appropriate.

### Ethical note

Data collection was observational and adhered to the International Primatological Society’s Code of Best Practice for Field Primatology (MacKinnon et al. 2014). All applicable international, national, and institutional guidelines for the care of animals were followed.

Research was conducted under approval by the Uganda Wildlife Authority and the Uganda National Council for Science and Technology. All work was in accordance with the ethical standards of the Budongo Conservation Field Station at which the study was conducted.

### Data collection

Researchers and a team of field assistants followed chimpanzees daily (Waibira: from 06:00 to 18:00; Sonso: from 07:00 to 16:30). Long-term data collection included continuous focal individual activity and party composition taken on a 15-min scan basis. In addition, all unusual events or otherwise remarkable behaviour were recorded in detail in logbooks for each community, including births, deaths, and associated descriptions of behaviour (Sonso: since 1993; Waibira: since 2011).

In addition to long-term records, AS, PF, EF, CF, DT and CH together with field-assistants SA, JA, GA, BC, and GM of the Budongo Conservation Field Station observed the extended carry events we report. KET and UP are typically comfortable with the presence of human observers; however, following the death of KETs infant we avoided selecting her as a focal individual because we observed behavioural cues known (or assumed) to be associated with greater than typical social and physiological stress (e.g., self-scratching and vigilance) in her interactions with other chimpanzees and we did not want our extended presence to further impact these. Observations of her behaviour were taken on an ad libitum basis whenever she joined the party of chimpanzees that included a focal individual, but we made an effort to locate and observe her for a brief period of time each day to obtain regular updates on her and her infant’s state of decomposition. During the births and deaths of UP’s infants, regular research practices had been adjusted due to the Covid19 pandemic. In 2020 activities were restricted to shorter hours of observation (7:30 to 13:00) and limited to CH and the permanent field staff, who focused primarily on health monitoring of the chimpanzees during this period; in late 2021, at the time of UP’s second extended infant carry, restricted research activities had resumed. Researchers and field staff opportunistically noted any unusual behaviour exhibited. Particular attention was given to how the bodies, and in one event an object that we suggest may have been a potential substitute for the dead infant, were transported, the response of nearby individuals to the mother or the carcass, the interactions of the mother with the corpse, and the state of the corpse. We were not able to collect physiological samples from either corpse to perform laboratory analyses on the cause of death, nor we were able to retrieve either body for autopsy.

## RESULTS

Over a combined 40-year period of observations (30 years Sonso, 10 years Waibira) a total of 191 births were recorded. Of these, 68 (36%) died in infancy (≤ 5 years) offering opportunities for their mothers to carry the infant’s corpse post-mortem. We found no evidence for seasonality as deaths (with a confirmed observation month, n=59) occurred throughout the year (Jan n=5; Feb n=2; Mar n=2; Apr n=3; May n=1; Jun n=3; Jul n=9; Aug n=6; Sep n=10; Oct n=6; Nov n=9; Dec n=3). Of the 68 infant deaths, we excluded three that died together with their mothers and 12 because they were partially dismembered or cannibalised during infanticides. Of the remaining 53 cases, for 46 (87%) we were able to estimate the infant’s age at death (±1 month). The majority (n=25; 54%) died within the first month, 17 (37%) at 1-month to 1-year old; 3 (7%) at 1- to 3-years old, and one (2%) at 3-to 5-years old.

We observed 12 carries of dead infants by their mothers (Table 1), 23% of observed opportunities (n=53). To be included as a case of dead infant carrying we required that the mother be seen with the infant the day after death was estimated to have occurred. In nine cases the minimum carry length observed was 1-3 days, in three cases we observed a longer minimum carry of n=18, n=56, and n= 89 days. These are described in more detail below. The 12 carries occurred in both primiparous (n=1) and multi-parous females (n=11), including a seventh born infant. However, these observations are likely an under-estimate of the frequency of dead infant carrying behaviour in Budongo mothers. In 36 instances the mother reappeared alone and could have carried for an unknown period prior to this. In total there were 29 cases where the mother and dead infant were seen together, of these 12 included a death with the mother or infanticide with cannibalism. Of the remaining 17 cases, 12 showed carrying of the dead-infant, a rate of 71%. Of the five cases where the mother was observed with the dead infant but did not carry it, all were infanticides (without cannibalism). Finally, four out of eight mothers (ML, JN, KU, UP) were observed carrying the dead bodies of two of their infants (Table 1), one carried for the same duration and three carried for longer durations on the second occasion.

**Table 1.** Carrying of dead infants by Budongo chimpanzee mothers: mother-infant dyads (with the mother first), mother’s age, parity (indicated as multi-parous (multi) or primi-parous (primi)), infant’s age, cause of death, duration of carrying, community (Sonso or Waibira), and date of infant death.

| Mother-infant | Age mother (years) | Parity | Age infant (months) | Cause of death | Minimum carrying duration (days) | Community | Date of infant death |
| --- | --- | --- | --- | --- | --- | --- | --- |
| KG-KG2 | 21 ±3 | multi | 0.03 | still birth (suspected) | 2 | Sonso | 09.09.1997 |
| ML-ML2 | 26 ±3 | multi | 0.03 | still birth (suspected) | 1 | Sonso | 08.07.2002 |
| JN-JN2 | 21 ±1 | multi | 0.5 | unknown | 1 | Sonso | 24.01.2005 |
| ZM-ZM6 | 41 ±5 | multi | 0.4 | unknown | 1 | Sonso | 19.03.2009 |
| JN-JN4 | 28 ±1 | multi | 0.25 | infanticide | 3 | Sonso | 10.11.2012 |
| KU-KU5 | 34 ±3 | multi | 0.1 | infanticide (suspected) | 2 | Sonso | 18.02.2013 |
| KL-KL7 | 34 ±1 | multi | 0.12 | infanticide | 2 | Sonso | 30.07.2013 |
| ML-ML5 | 39 ±5 | multi | 0.06 | infanticide (suspected) | 3 | Sonso | 17.11.2014 |
| KET-KYO | 20 ±1 | primi | 25 | respiratory infection (inferred) | 18 | Waibira | 06.01.2018 |
| KU-KU7 | 40 ±3 | multi | 0.15 | unknown | 2 | Sonso | 26.11.2019 |
| UP-UP3 | 21 ±1 | multi | 0.25 | unknown | 56 | Sonso | 23.09.2020 |
| UP-UP4 | 22 ±1 | multi | 0.25 | unknown | 89 | Sonso | 21.08.2021 |

We found no evidence for a clear effect of seasonality on the onset of carrying behaviour, but our sample size is small (see Fig. 1). The Budongo Forest shows two marked periods of intense rainfall (March – May, September – November), an intense drier season (December – February) and a light drier season (June – August). Carrying of dead infants was as likely to occur in the Wet (n=6) or Dry (n=6) seasons (X^2^ = 0.00, p=1.000).

**Fig. 1.**
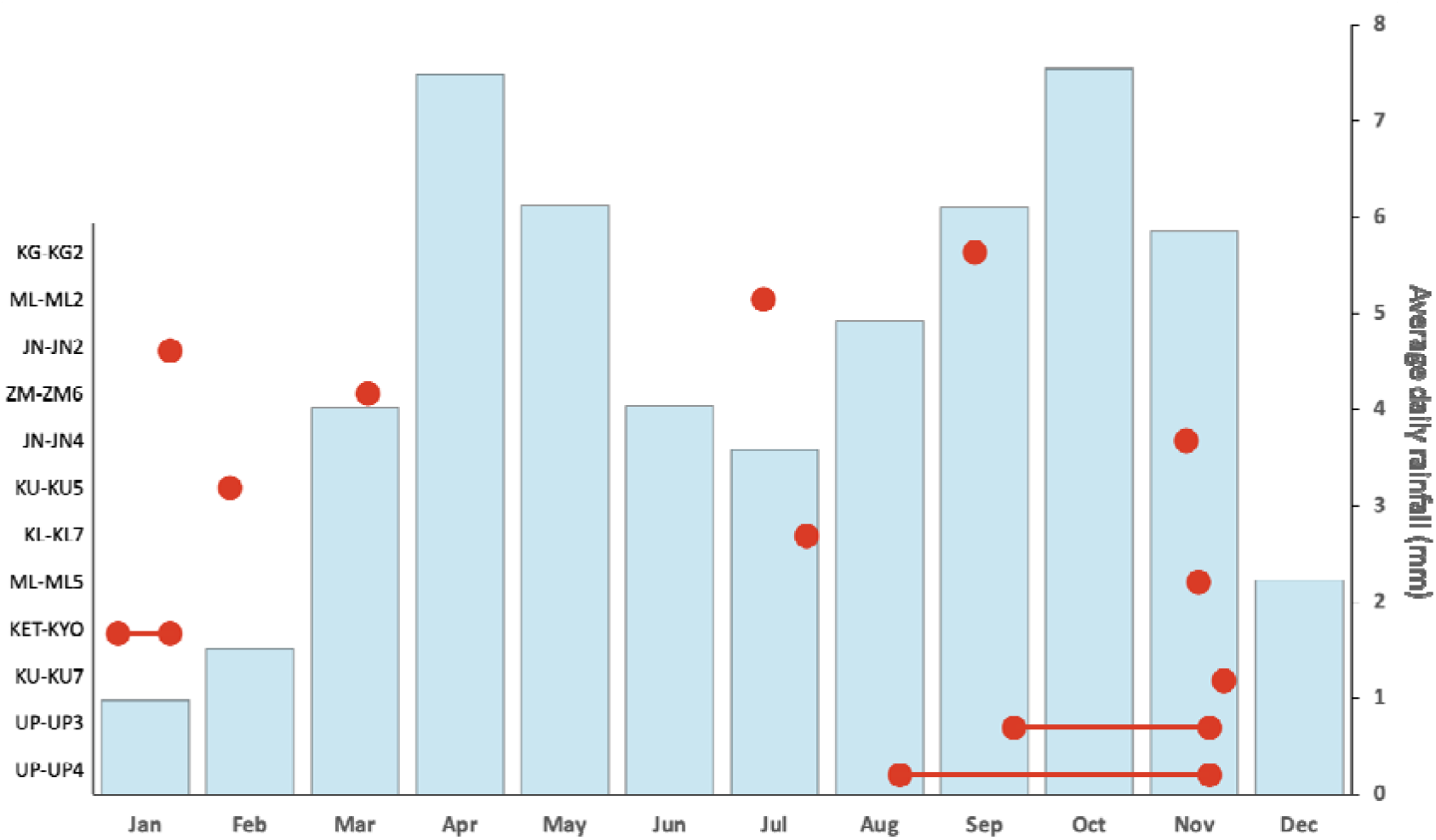
Dead infant carrying occurrences for each mother-infant pairs relative to the average daily rainfall (data extracted from Budongo Conservation Field Station long-term records 1993-2018). Single red dots represent occurrences of carrying that lasted between 1 and 3 days. Red dots connected with a line represent the approximated duration of extended carrying, with the dots representing the start and end of carrying.

**Fig. 2.**
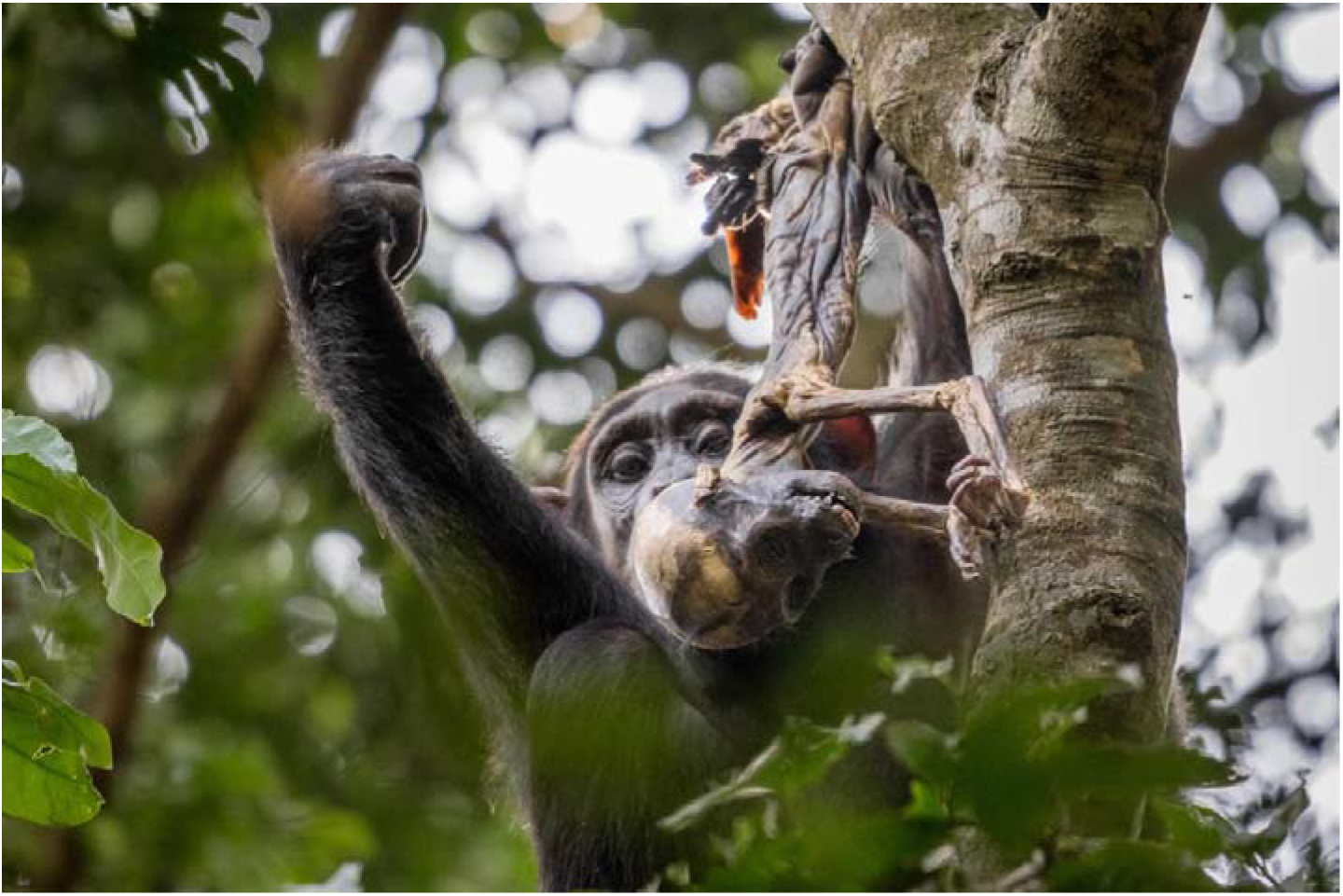
KET holding KYO’s mummified body while sitting on a tree (picture taken by co-author on the 21.01.2018)

**Fig. 3.**
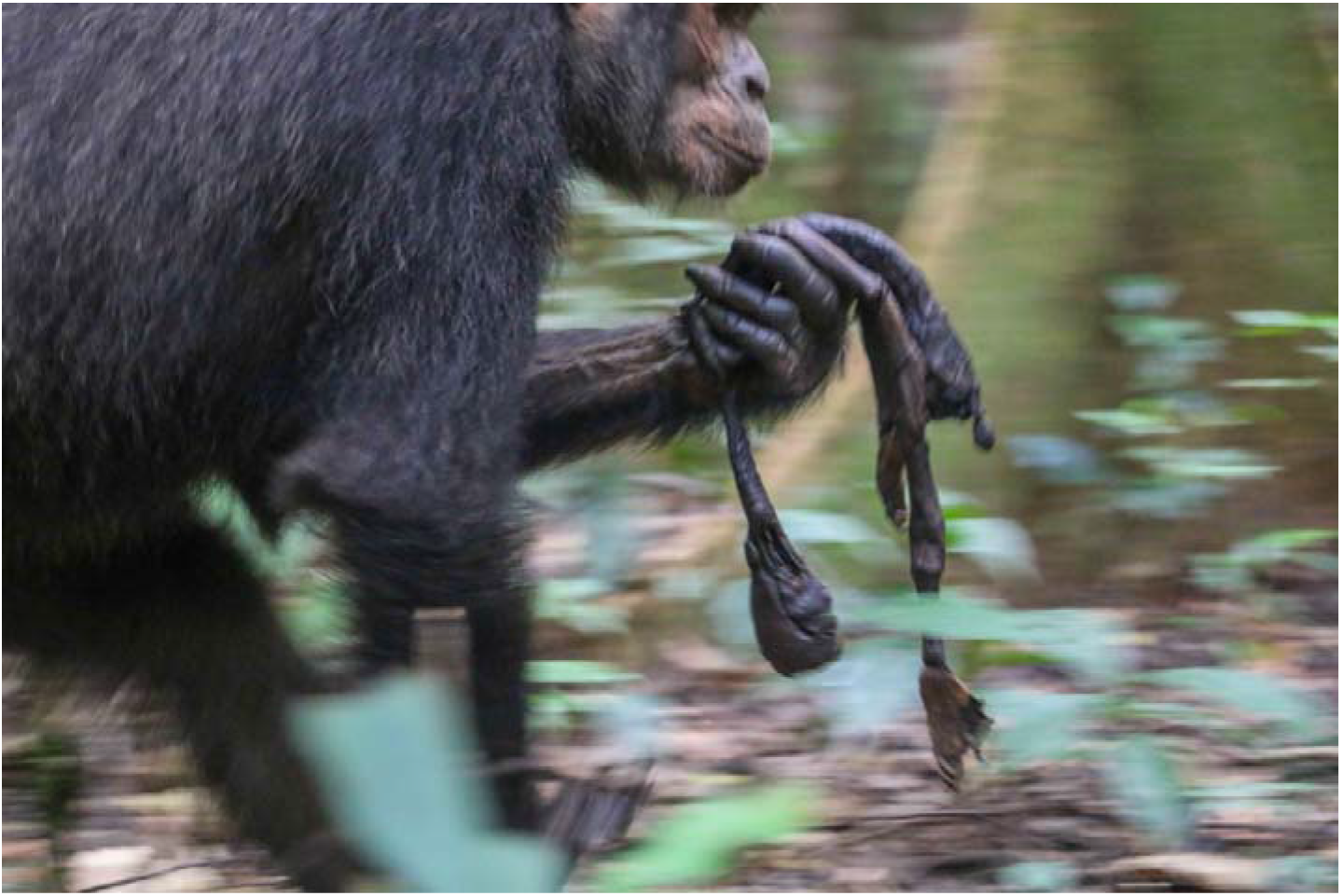
UP carrying UP4’s mummified body in her hand while traveling on the ground (picture taken by co-author on the 30.08.2021)

We found a possible effect of infant age. Our sample size of older infants was very small: of the 46 opportunities to carry described above, we only observed four infants who died between 1-5 years old, and a single case (25%) of carrying (aged 2-years, described below as the extended carry by KET of KYO). We found a similar frequency of carries (n=11), given opportunities to carry (n=42) of infants under one year old. However, within this larger sample of younger infants aged 1-12 months, we found that only very young infants (aged under 1-month) were carried. When considering the number of opportunities to carry, infants aged under 1-month (11 of 25 cases, 44%) were more likely to be carried than infants aged 1-12 months (0 of 17 cases; Fisher’s exact test p=0.010).

### Extended dead infant carrying

A detailed description and videos of the observations are available in the Supplementary Materials. Here we provide a summary of the key information.

### Observation 1: KET, extended dead infant carrying in Waibira

KET’s first born infant KYO was last seen alive on the 6th January 2018, aged 25 months. On the 7th January 2018 KET was observed carrying KYO who appeared lifeless. Other chimpanzees were present and were apparently aware of her arrival with the infant but showed no atypical reactions to KET or to the corpse. Based on the fact that most chimpanzees exhibited signs of respiratory infection during that period and given the infant had shown no other sign of illness, the likely cause of death was inferred to be respiratory infection. During the first day KET was observed scratching herself repeatedly before approaching a water area and when sitting close to a sub-adult male. These scratches appeared to be stress-related (fast and repeated, and not accompanied by grooming or response waiting). On several occasions she moved her hand over the dead body apparently to chase away the flies. Other than this, during the entire 18-day period, she was never observed to provide any direct maternal care (grooming, inspecting, touching, or peering) other than carrying, and regularly left the body at short distances (up to 5 m) without visually monitoring it. She did not stop others from approaching herself or the dead infant. When moving or feeding in a tree, the dead body was usually (15 out of 16 observations) placed in her right leg pocket, when on the ground she carried the body in her hand or arm (see Online Resources). On one occasion a nulliparous young adult female (MON) was observed to briefly carry the corpse in one hand while KET followed her. Across the 18-days KYO’s corpse decomposed, initially increasing in smell. By the 4th day, no hair remained on the body. By the 9th day, the body looked “dried”. By the 10th day the pungent smell and number of flies decreased. It is likely that at this stage the body was completely mummified. No other chimpanzees responded noticeably to either the smell or the flies. On the last day of observed carrying, the body was still intact with only eyes missing and one deformed ear.

### Observation 2: UP, extended dead infant and subsequent object carrying in Sonso

UP was first seen on the 25th September 2020 with an apparently recently dead infant (UP3), estimated to be 1-week old. Her two previous infants were killed by intra-community infanticide when under 2-weeks old. While some immature individuals (< 10 years old) inspected the carcass, no others did, and an adult male showed no interest even while grooming UP. The infant’s corpse had started to dry out, but had a noticeable smell and flies, and was assumed to have died several days earlier. UP was observed on the 4th and 26th October and the 8th and 19th November, carrying the corpse on all occasions. She held it in her hand when on the ground and moved it to a leg pocket when climbing or moving in trees. By the 8th November the corpse appeared fully mummified. UP was last seen with the corpse on the 19th November, a minimum carry duration of 56-days (although likely several days longer given corpse appeared partially dry on first observation). She was next seen on the 23rd November. When approaching a water hole at the base of a tree, she carried a twig ∼50 cm long in her mouth. As she reached the hole she transferred the twig to her hand and left leg pocket, drank, and then returned it to her mouth (see Online Resources). She continued to carry the twig throughout the morning, including while climbing large trees, and when patrolling with the group over several kilometres. She was seen on the 24th November, and 3rd and 4th December, and was again observed to be carrying a similar twig consistently. She was not seen to put it down on the ground. She was seen briefly on the 5th December, but it was not clear if she had a twig with her, and when she was next observed in the new year (30th January) she no longer carried anything.

### Observation 3: UP, 2nd extended dead infant carrying in Sonso

UP was first seen on the 28th August 2021 with another dead infant (UP4), estimated to be 1-week old. While we did not see her with a live infant, and it is thus at least possible that the infant was not hers, we believe this to be very unlikely. UP was last observed in maximal oestrus approximately 8 months prior to being observed with the dead infant, and all other Sonso females were either confirmed to be pregnant, had unweaned infants at the time, had been observed cycling regularly in the months prior to the observation, or were considered to be post-reproductive. Finally, we know of no other reports of extended carrying of non-kin dead infants. The infant’s corpse had started to dry out but had a strong smell and on the 30th August flies could be seen hovering around it. Given that the corpse still had a strong smell but was already partially dried, it was assumed to have died at least one week prior. The cause of death was unclear; however, UP was observed with wounds on her head and on her left arm. UP was observed using three main carrying styles when traveling on the ground or moving in trees. She either carried the corpse in one hand (typically left one), in the mouth, or in one leg pocket (typically left one) (see Online Resources). When resting, she placed the corpse on her lap, in a leg pocket, held it in one hand, or placed it on the ground close to her. UP was not observed providing direct maternal care (e.g., grooming, inspecting, or peering) to the corpse, though she was observed moving her hand around the dead body to chase away flies on several occasions. On one instance UP was victim of aggression from other females during which she dropped the corpse, and then followed the group when traveling and left the corpse behind. Soon after, she was observed returning to retrieve the corpse. We observed a juvenile male orphan (KJ) following her and peering close to the corpse on a few occasions. No other individual was observed taking interest in or showing response to the corpse.

Throughout the observation period, UP was often seen in large groups and regularly socialising with adult males (e.g., grooming). On this occasion there were no observations of object carrying. UP was seen carrying the corpse on the 28th, 30th and 31st August; the 2nd, 6th, 8th, 9th, 11th, 13th, 14th, 16th, 18th-25th and 29th September; the 7th, 12th, 13th, 15th-21st, 26th, 28th October; and the 3rd, 10th, 14th and 17th November. On the 18th November she was seen without the corpse and had resumed her sexual cycle (with visible sexual swelling) for the first time since the last pregnancy.

## DISCUSSION

We compiled over 40-years of long-term data on chimpanzee mothers from the Budongo forest that had lost their infants but continued carrying them for days. Dead-infant carrying was practiced by both parous and primiparous mothers with both new-born and older infants. If we consider only those in which we were able to observe the mother immediately after the infant’s death, it occurred in at least a fifth of cases. However, this value is very likely an under-estimate of the frequency with which bereaved Budongo female chimpanzees carry their infants. The Sonso community, in which we made most of our observations, experiences periodic high levels of infanticide (Lowe et al. 2020). These infanticide cases are often accompanied by some level of cannibalism or dismemberment, and/or the infant is taken from the mother (Lowe et al. 2019), which may limit or shorten mothers’ opportunities to carry (Fernández-Fueyo et al. 2021; Gonçalves and Carvalho 2019). In many cases the mother was simply not seen in the days following her infant’s death. In the limited number of cases where the mother was seen in the days following the infant’s death and where the death was not an infanticide with cannibalism of the corpse, almost three quarters of cases involved carrying of the dead infant.

We found no seasonal effect on dead infant carrying, but we do show a possible age effect: amongst infants who died at under 1-year old, very young infants were more likely to be carried. Our observations also support the suggestion that infants may be less likely to be carried following traumatic death (Fernández-Fueyo et al. 2021). Most cases of infant carrying were relatively short (minimum confirmed carry length of a few days); however, we also reported three prolonged cases of extended infant carrying. Our observations suggest that these mothers, despite the evidence of irreversible loss including absence of any resemblance to living infants, continued to experience a strong attachment to their deceased infants.

Neither female had any other living offspring and one, after eventually abandoning her dead infant after 56 days, carried an object (a twig) for at least another two weeks. In the three cases of prolonged carrying that we report, dead-infant carrying was not accompanied by other forms of maternal care, such as grooming or other forms of maternal attention or interactions (e.g., Matsuzawa 1997; Biro 2011), suggesting that the two mothers had become aware of the biological facts. Both mothers were forced to use atypical modes of infant carrying, including mouth carrying, more typically used for objects (Lonsdorf et al. 2020). Overall, these data suggest that the ‘unawareness hypothesis’ is unable to fully explain chimpanzee behaviour towards dead conspecifics.

As neither mother appeared to inspect or interact with the infant beyond carrying, our observations do not support the ‘learning about death’ hypothesis. Further support against the ‘learning about death’ hypothesis is provided by the mothers who carried their dead infants on more than on occasion: of the four mothers, three carried for longer on the second occasion (one for the same amount of time). When we extracted the same data from Lonsdorf et al. (2020, n=6), Biro (2011, n=1), and Hanamura et al. (2015, n=2), this pattern appeared repeated: six of the nine cases described were longer on subsequent carries. Recurring and prolonged carrying behaviours seem to indicate that mothers are not unaware of death.

While both KET and UP were inexperienced mothers (primiparous and parous but all offspring killed at under a month old respectively), 10 of the 11 mothers who carried their dead infants were parous and two of these cases were with sixth and seventh born infants. Cases of extended carrying by parous mothers in other groups also suggest limited support for the ‘learning to mother hypothesis’ (Matsuzawa 1997; Biro et al. 2010; Biro 2011; Lonsdorf et al. 2020). However, in line with the fact that younger primate mothers are more likely to carry dead infants (Fernández-Fueyo et al. 2021), our three observations support the suggestion that rare instances of extended carrying across several months might be more frequent in young mothers. Nevertheless, repeated prolonged carries do not support the suggestion that this is due to inexperience and, given that no clear pattern emerges when considering observations across sites (e.g., Hanamura et al. 2015; Lonsdorf et al. 2020) and that sample sizes of chimpanzee mothers remain very small, these rare instances could also reflect individual differences.

Because we did not collect any hormonal data to assess the levels of stress associated with dead infant carrying, we were unable to evaluate the ‘grief-management hypothesis’. However, a recent study by Girard-Buttoz and colleagues (2021) reported elevated cortisol levels in infant chimpanzees who lost their mothers, supporting the notion that disruption of the mother-infant bond leads to elevated stress. Similarly, female baboons experience high levels of glucocorticoids when losing an ally to predation and in periods of infanticidal attacks (Engh et al. 2006a, b). Given the bond chimpanzee mothers share with their infants is among their most significant (Pusey 1983; Lonsdorf and Ross 2012; Stanton et al. 2017), we expect mothers to experience elevated stress levels following the death of their infant.

One of the three prolonged carries took place during the peak of the driest season, whereas the other two took place during the wettest season, and we found no effect of seasonality in our wider data. Rather than mummification being the result of favourable climactic conditions, it is possible mummification was observed because the extended carrying durations allowed for it (see also Biro 2011). Recent explorations of several large datasets also found no support for a ‘climate hypothesis’ (Das et al. 2019; Lonsdorf et al. 2020; Fernández-Fueyo et al. 2021). Of the three extended carries two infants were new-born, while the other was 2-years old. The longest carry reported was for a new-born, however, other new-born infants were carried for short periods of just a few days. One of the three extended carries terminated with the resumption of the mother’s reproductive cycles. While our observation of carrying being more likely in very young infants (under 1-month) than in infants aged 1-12 months, as well as the extended carrying by UP of her two young new-borns would fit the pattern proposed for the ‘post parturient hypothesis’; however, the extended carrying by KET of her 2-year-old infant does not. Previous studies suggest that while post parturient effects may contribute to this behaviour, they cannot explain extended carrying alone (Watson and Matsuzawa 2018; Masi 2020). However, hormonal data are needed to investigate this hypothesis effectively. While KET’s case provides support for the ‘maternal-bond strength hypothesis’, UP’s cases provide counterevidence. However, given that the latter were UP’s third and fourth infants in a five-year period, the first two having been killed at under a month old in within-community infanticides, it is difficult to assess the nature of her bond with these infants. Soon after dropping the corpse, UP was observed carrying a twig for several days, which we suggest may have been used as a substitute for her dead infant’s body. This unusual behaviour together with the even more prolonged second carry suggest she had a particularly strong motivation to carry. Our observations (see Table 2 for a summary), combined with the fact that all recorded instances of carrying in our dataset concern infants who died before weaning age, seem to indicate that maternal behaviours, which are not limited to maternal care, and the bond between the mother and the offspring likely play an important role in dead infant carrying (Fernández-Fueyo et al. 2021).

**Table 2.** Hypotheses, predictions, and supporting evidence from the present study for dead infant carrying

| Hypothesis | Prediction | Study support |
| --- | --- | --- |
| Unawareness | Dead infants are treated as alive. | Unlikely |
| Post-parturition | Young infants are carried for longer. | Mixed |
| Learning about death | Mothers inspect and check state of infant. | Unlikely |
| Grief-management | Stress levels are lower in mothers carrying dead infants. | N/A |
| Learning-to-mother | Primiparous mothers carry dead infants more often/for longer. | Unlikely |
| Maternal-bond strength | Strongly bonded and intermediate/old infants are carried for longer. | Mixed |

Within our observations of dead infant carrying, there were two examples of particularly rare behaviour: KET allowing MON to briefly carry KYO, and UP’s twig carrying. Carrying the infant of others is an unusual behaviour in East African chimpanzees. It has been observed on rare occasions in Budongo: two adult males snatched new born infants and carried them (still alive) for at least two days (in one case the male continued to carry the infant for a further two days after its death; Notman and Munn 2003; unpublished long-term data), and a daughter was observed carrying what was suspected to be her mother’s new infant for several days (unpublished long-term data). It is possible that KET tolerated MON’s brief carry because they may share a close bond, but another explanation is that her own bond with the infant’s body had perhaps decreased by the 13th day.

We are not aware of any other reports of primate mothers carrying substitute objects following their infant’s death and we are cautious about interpreting this unique and unusual observation. While chimpanzees may carry objects for many reasons, a number of features suggest that the observation of UP carrying a twig was related to infant carrying. Prior to the incident, neither UP nor any other adult Budongo chimpanzees had ever been observed to carry non-food objects between locations. They are notoriously non-stick-tool users (Whiten et al. 1999; Gruber et al. 2009, 2011; Gruber 2016). Chimpanzees have been reported at several sites to engage in ‘doll’ play, where substitute objects, including logs and sticks, are carried as if they were a young infant (Matsuzawa 1997; Kahlenberg and Wrangham 2010). This behaviour typically peaks in juveniles and is more frequent in females and while it is observed in some adult females it ceased once they became mothers (Kahlenberg and Wrangham 2010). The description of log doll use in Bossou is of particular interest here, as it was carried by a juvenile female during the period that her mother was carrying her sick infant sister, who subsequently died and whose body was also then carried (Matsuzawa 1997). UP’s behaviour was observed multiple times over several weeks and, unlike the descriptions of other ‘dolls’, she was not seen interacting with the object, treating it instead in the same way as she had her infant’s corpse. Thus, object-carrying may also be associated with the loss of an infant in bereaved chimpanzee mothers. In humans, the use of transitional objects (e.g., dolls or objects associated with the deceased) has been suggested to function as a coping mechanism following a bereavement (Graham et al. 1987; Lister et al. 2008). A similar suggestion has been made for beluga whales where both wild (Smith and Sleno 1986) and captive (Kilborn 1994) individuals have been seen to carry inanimate objects, apparently as ‘surrogates’. The captive whale carried a buoy followed the removal of her dead calf immediately after birth (Kilborn 1994), and in the wild observations included carrying of planks and netting (Smith and Sleno 1986). Further observations are necessary to validate the use of object-carrying following the death of an infant as a coping mechanism in primates.

To sum up, our observations are consistent with previous observations that chimpanzee mothers respond to the death of their infants with carrying behaviour across communities. Furthermore, our observations support the argument that these mothers act as if they are aware of the loss but continue to display a strong attachment to the bodies of their infants and may be affected by psychological processes akin to human grieving. Nevertheless, more detailed hormonal data are needed for a test of this potential mechanism. A combination of ecological conditions favouring mummification, and social factors, such as the strong bond shared between mothers and their infants, may explain the three particularly extended carries by Budongo chimpanzees. While we did not observe other indications of maternal care in these cases, we are cautious about interpreting this as a wider absence in Budongo mothers. Mothers’ pattern of behavioural responses to death may be individually specific and nuanced, resulting from a combination of physical, ecological, and psychological factors, and more observations are needed to generalise at the population or species level. Our interpretations are limited by the small number of observations and the multitude of possible influential factors to consider. We encourage researchers and long-term field sites to continue to report the rare behaviours observed in different populations, for example by contributing to open-access databases such as ‘ThanatoBase’ (http://thanatobase.mystrikingly.com), to allow a richer exploration and more robust hypothesis testing of non-human primates’ reaction to death through data sharing and collaborations across sites.

## Supporting information

Online Resource 1

Online Resource 2

Online Resource 3

Online Resource 4

Online Resource 5

Online Resource 6

Online Resource 7

