## Supplementary materials for "Dead infant carrying by chimpanzee mothers in the Budongo Forest"

^2^ Department of Comparative Cognition, Institute of Biology, University of Neuchâtel, Neuchâtel, Switzerland

*Detailed reports of extended dead infant carrying*

**Observation 1: KET, extended dead infant carrying in Waibira**

KET’s first born infant KYO was last seen alive on the 6th of January 2018, aged 25 months. On the 7th of January 2018 at around 10:30am KET was observed carrying KYO who appeared lifeless while approaching a water source. Other chimpanzees were present and were apparently aware of her arrival with the infant, but they showed no atypical reactions to KET or the carcass. The body of the infant was held and carried by KET ventrally with one hand, and there were no signs of movement from KYO. We suspected KYO was dead when KET left the body a few meters behind her on the ground without monitoring as she moved to drink. While infanticide is relatively common in Budongo (Lowe et al. 2019, 2020), the body showed no external signs of attacks or other wounds. Based on the fact that most chimpanzees in this community exhibited signs of respiratory infection (coughing, sneezing, and mucus) during that period of time and that the infant had shown no other sign of illness, we suspect that the most likely cause of death was as a direct or indirect result of the respiratory infection. At the time of KYO’s death, it was unclear whether KET had clear symptoms of infection (absence of coughing and sneezing, but she picked mucus from her nose), but 10 days after the death of her infant she was recorded with a serious cough. Other chimpanzees in this community, including a mother of dependant offspring (SHU and her new-born (died), SPN, SDT (survived)) were recorded missing over the same period of time. Lethal respiratory outbreaks have become more regular in Ugandan chimpanzee populations in recent years (Scully et al. 2018), and have occurred regularly in other chimpanzee populations too (e.g. Köndgen et al. 2008). However, KYO had consistently appeared small and under-developed for her age. She was more similar in size to 1-year old infants and appeared to be less independent than her age-mates (AS, CH personal observations).

*KET behaviour*

During the first day of observation, KET was observed scratching herself repeatedly before approaching the water area and when sitting close to a conspecific sub-adult male. These scratches appeared to be a sign of arousal, rather than for hygiene or gestural communication, as they were not accompanied by grooming or response waiting and occurred with increased frequency, repetition, and speed. While resting on the ground, KET left the dead body adjacent to her and no further than 5 meters from her, but she did not regularly visually monitor it. For example, while approaching a water source to drink she left the body 5 meters behind her and did not monitor its state, even where others, including adult males, approached within 5-10 meters. This is in contrast to what a mother with a living infant of KYO’s age would typically do when physically distant. On several occasions she moved her hand over the dead body apparently to chase away the flies (Online Resource 3). During the entire 18-day period she was never observed to groom the body, nor to inspect it from a close distance or for an extended period. Furthermore, when leaving it on the ground, she did not regularly visually monitor the body. We did not record any other indicators of direct maternal care, such as grooming, inspecting, touching, or peering, even though she carried the dead body for an extended period of time. KET did not stop others from interacting or approaching herself or the dead infant.

*KET carrying style*

Over the 18-day period KET was observed transporting the dead body in different ways. Based on data from video recordings, we identified clear instances of transport or carrying (n=35) across resting (12), feeding (6), and moving (17) contexts both on the ground (19) and up in trees (16). During resting periods, she would usually leave the dead body on the ground at around 1 meter from her, but when in a tree or when resting for brief intervals, she carried the corpse in the same fashion she used during traveling. When moving or feeding in a tree, the dead body was usually (15 out of 16 times) placed in her right leg pocket with the head of the carcass facing outward, though she was also observed supporting it over her right forearm. In contrast, when on the ground she carried the body in 3 different manners: in her right hand (5 out of 10 times), on her right forearm (3/10) (Online Resource 1), and in her left hand (2/10). During most carrying instances, the head of the infant was positioned facing outward her own body (legs and lower body facing inward) (Online Resource 2). On one occasion a nulliparous young adult female (MON) was observed briefly carrying the corpse in one hand while KET followed her.

*State of corpse*

We recorded the level of decomposition of KYO’s dead body from the first day of observation. The smell of rotting flesh increased rapidly and after a couple of days her presence could be smelled before she was in sight. After 3 days, the corpse had changed drastically, with hair absent in all areas of the body except the head. 4 days after there was no more hair on the head. On the 9th day, the body had significantly reduced in size and acquired a “dried” and “pale” look. The pungent smell decreased slightly and became stable after approximately 10 days. The number of flies seemed to decrease around the same time. It is likely that at this stage the body was completely mummified. No other chimpanzees responded noticeably to either the smell or the flies. On the last day of observed carrying, the body was still intact with only eyes missing and one deformed ear.

Observation 2: UP, extended dead infant and subsequent object carrying in Sonso

UP was first seen on the 25th September 2020 with an apparently recently dead infant (UP3), estimated to be 1-week old. She had not been seen since the 5th July 2020. Her two previous infants were killed by within-community infanticide (UP1 24th July 2017 at 2-days old; UP2 19th July 2018 at an estimated 2-weeks old). At the time she was observed she was with two other individuals, shortly afterwards two adult males joined the party and displayed towards her. One of them (FK) approached her and started to groom her but showed no apparent interest in the carcass. A subadult female (HR) whose family had joined the party approached her and inspected the dead infant on several occasions, as did a 4- and 6-year-old infant. The infant’s corpse had started to dry out, but still had a noticeable smell and fly activity, and so was assumed to have died several days earlier.

*Carrying the dead infant*

UP was next observed on the 4th October carrying the corpse. She held it in her hand most of the time and moved it to a leg pocket when climbing or moving in trees. She was again observed to carry the corpse on the 26th October, and the 8th and 19th of November. When seen, she moved with the main group and socialised with others. By the 8th November the corpse appeared fully mummified, it was completely dry, had lost a lot of hair, and no longer noticeably smelled. UP was last seen with the corpse on the 19th November, a minimum carry of 56-days (although likely at least several days longer given corpse appeared partially dry on first observation).

*Object carrying*

On the 23rd November at 9:40 UP was observed to approach a water hole at the base of a tree. She carried a twig ~50 cm long with a few wilted leaves on it in her mouth. As she reached the hole she transferred the twig to her hand and to her left leg pocket, drank, and then returned it to her mouth before walking away (Online Resource 7). She continued to carry the twig throughout the day, including while climbing into a large fig (*Ficus mucuso)* tree, and when patrolling with the group over several kilometres of their Eastern territorial boundary. At 13:00 she was seen without the twig. While she moved with the group she sat on the periphery and was not observed to interact socially, including direct requests to play from a juvenile. On the 24th November she was again seen, and observed to carry a similar twig throughout the morning, but again had dropped it by early afternoon. She was seen on the 3rd and 4th of December, and again observed to carry a twig (~25 cm) with a few yellowed leaves on it in her mouth consistently throughout the day when travelling or climbing, and while feeding she would hold it in her leg pocket. She was not seen to put it down on the ground. She was seen briefly on the 5th December, but it was not clear if she had a twig with her, and when she was observed again in the new year she no longer carried anything.

**Observation 3: UP, 2nd extended dead infant carrying in Sonso**

UP was first seen at 07:26 on the 28th August 2021 carrying a dead infant. UP was last seen on the 24th of July before the observation reported here. When observed for the first time with her dead infant, UP was feeding with eight other group members from two families (two adult mothers with two infants, two juveniles, and two sub-adults). The group was observed in a large fig tree (*Ficus varifolia*). The corpse of the infant had started to dry out and most of the flesh appeared to have decayed leaving behind long thin remains. Due to the decomposition of the corpse, the cause of death could not be ascertained. The corpse had a strong smell and flies nearby, which UP occasionally swatted away with her hands. Given the time last seen, and the size and status of the corpse we estimate the infant to have been under 2-weeks old at time of death, and to have been dead for at least one week, giving an estimated date of birth of around the 7th August ± one week.

UP was observed for the second time on the 30th of August at approx. 07:50. A juvenile male (KJ) and an adult male (MS) were feeding on figs (*Ficus varifolia*) when UP approached them carrying the corpse of the dead infant. UP had the corpse in her mouth while she moved in the tree and the corpse’s face was still distinguishable (Online Resource 4). At around 08:05 UP, MS, and KJ descended from the tree. After travelling, the group stopped and KJ appeared to peer twice at the corpse from a close distance (<0.5 m). At 08:19 UP groomed MS and KJ was seen peering, though it was unclear if at grooming or at the corpse. When they resumed traveling, KJ inspected the ground where UP had sat from a very close distance. At 08:24 the group stopped and sat along a log tree. KJ peered twice at the corpse. At 08:36 UP approached an adult male (PS) who had displayed when the two groups merged. PS descended the tree and mutually groomed with UP for approx. one min. At 08:40 another adult male (MS) together with the alpha female (and his mother, NB) joined UP’s group and MS displayed at UP chasing her up a tree. The males and KJ left the group. Another male (KT) joined the group at 08:56 and approached the tree where UP was resting. After one min, she descended and started to groom KT before he moved ~3 m away. UP approached KT again at 09:05 and groomed him, but he quickly turned and again moved ~3m away. At 09.10 UP followed KT and they mutually groomed for approx. five mins before KT left. While UP was self-grooming, she touched her swelling and smelled her hand, and then appeared to swat flies away. We noticed she had a wound on her head and was licking a wound on her left arm, possibly from a recent attack. At the time, the corpse smelled strongly (very noticeable at ~10 m distance) but the other individuals did not seem to react to it. At 09:53 UP travelled alone. She then moved to a feeding tree (*Ficus exasperata*) where she joined other individuals. We observed PS grooming UP. At around 10:10 PS moved away and KJ moved within 3 m from UP, but she quickly moved away and fed on fruits. The whole group moved out of the tree soon afterwards. When we found them resting, we observed an adult female (KL) inspecting UP’s swelling with her fingers and then smelling them. UP then approached KT and they alternately groomed one another. KJ peered at UP grooming KT while a sub-adult male (KC) moved away when UP approached. When UP approached KC, he began grooming UP while KJ peered with his head very close to the corpse (<0.5 m) (Online Resource 6). When KC and KJ left, UP remained behind placing the carcass on the ground while self-grooming. At around 10:53 UP approached MS and groomed him and we noticed he turned his head away (the only potential response to the smell that we observed). After hearing calls from another party, UP vocalised in response (Online Resource 5), while MS and PS displayed and moved away.

On September 15th, a young primiparous female (KX) appeared with a newly born infant. While in the presence of males, KX attacked UP. UP did not show any behaviours that appeared to provoke KX. KX is a Sonso born female whose mother and maternal siblings (juvenile brother, and adult sister who also has her own infant) are currently in the community. UP did not show any apparent sign of stress after the event. On September 25th she was observed in a group with multiple individuals including three females, KA, KX, and KY (mother of KX and KA), all with their dependent offspring. KY and KX interacted agonistically with UP and produced loud vocalisations for ~ 1 min 20 s, which prompted her to drop the corpse. UP was seen without the carcass immediately afterwards. The group continued traveling while UP returned back to the previous location of the interaction. UP was seen later traveling with the corpse in her hand when she re-joined the group.

UP was seen carrying the corpse on: the 28th, 30th and 31st of August; the 2nd, 6th, 8th, 9th, 11th, 13th, 14th, 16th, 18th-25th and 29th of September; the 7th, 12th, 13th, 15th-21st, 26th, 28th of October; and the 3rd, 10th, 14th and 17th of November. On the 18th of November she was seen without the corpse and had resumed her sexual cycle (with visible sexual swelling) for the first time since the last pregnancy. She was again seen on the 20th November with sexual swelling and without the corpse. She was seen regularly over the following month and was not observed to carry a substitute object at any time. Overall, UP was often observed in large social groups throughout the observation period, including regularly socialising with mature males (grooming and traveling together).

*Carrying the dead infant*

UP was observed using three carrying styles when traveling or moving with the corpse across the observations: in her hand, in her mouth, and in her leg pocket. When she sat or rested, she placed the corpse on her lap, in her leg pocket, tucked under one arm, held it in her hand as if ‘folded’ (Online Resource 5), or placed it within a meter from her on the ground. When she travelled on the ground she often held it in left hand or carried it in her mouth, while when moving in a tree she carried it in her leg pocket or mouth (Online Resource 4).
